# Preclinical Development of a Vectorized Artificial miRNA Gene Therapy for Tauopathies

**DOI:** 10.1101/2025.10.12.681935

**Authors:** Irvin T. Garza, Brina Snyder, Sydni K. Holmes, Katherine M. Pearce, Krishanna Knight, Rachel M. Bailey

## Abstract

Tauopathies, including Alzheimer’s disease, are neurodegenerative disorders characterized by the accumulation of microtubule-associated protein tau, which is closely linked to cognitive decline. Reduction of tau is a potential and promising strategy for addressing tau-linked brain disorders. We report the development of a therapeutic approach using adeno-associated virus mediated delivery of an artificial microRNA targeting human tau. In a tauopathy mouse model, we demonstrate that a one-time intra-cisterna magna administration of vector resulted in reduced total tau, decreased pathological tau seeds, fewer tau inclusions, and amelioration of tau-related neuropathology. Notably, intervention at late disease stages, after onset of tau deposition and neurodegeneration, improved quality of life and extended survival. We further demonstrated the durability of therapeutic benefit and defined the minimally effective dose in tauopathy mice. These findings provide preclinical support for the advancement of a vectorized tau-lowering strategy as a disease-modifying approach for tauopathies and enable progression towards an investigational new drug application.

## INTRODUCTION

Microtubule-associated protein tau (MAPT, or tau) plays a critical role in maintaining neuronal cytoskeletal integrity^1,2^. During aging, tau can aberrantly dissociate from microtubules and misfold into toxic insoluble oligomeric species through mechanisms that remain incompletely understood. These oligomers act as seeds, promoting the nucleation and maturation of larger fibrillar aggregates known as neurofibrillary tangles (NFTs), which are a pathological hallmark of tauopathies^3–6^. Evidence supports that these seeds can move between cells and synapses in the brain and propagate the spread of pathological forms of tau along neural networks^5,7,8^. Tauopathies comprise a subset of neurodegenerative disorders characterized by the accumulation of tau aggregates as a primary or secondary pathological feature, and include frontotemporal dementia (FTD)^9^, progressive supranuclear palsy (PSP)^10^, chronic traumatic encephalopathy (CTE)^11^, and Alzheimer’s disease (AD)^12^. While antibodies targeting amyloid aggregates are approved for use in AD patients, these drugs incompletely alter tau protein accumulation and implore further therapeutic benefit.

Given that there is a close correlation between tau deposition, neurodegeneration, and functional decline^12,13^ therapeutic strategies aimed at reducing tau burden hold considerable promise. Approaches under investigation include targeting tau post-translational modifications^14–16^, immunotherapeutic modalities^17^, and antisense oligonucleotides (ASOs)^18^ designed to suppress tau translation. ASO-based therapies have demonstrated efficacy in mitigating neuropathology in disorders such as amyotrophic lateral sclerosis^19^ and tauopathies^18^. However, a limitation of ASOs is their relatively short half-life, necessitating repeated administration and imposing an additional burden on patient quality of life.

One strategy to address this limitation is to deliver the tau silencing agent using recombinant adeno-associated viral (rAAV) vectors. AAVs are promising delivery vehicles as they are small and relatively safe with hundreds of ongoing clinical trials, including the brain^20,21^. Although rAAV is constrained by a limited packaging capacity, it is well-suited for the delivery of small non-coding RNAs, including short hairpin RNAs (shRNAs), small interfering RNAs (siRNAs), and microRNAs (miRNAs)^22–24^. Artificial miRNAs (amiRNAs or amiRs) can be synthetically designed to make use of the endogenous RNAi cellular machinery to effectively silence genes of interest^23–27^. Accordingly, an rAAV-mediated delivery of a miRNA targeting *MAPT* transcripts could effectively inhibit the accumulation of pathogenic tau by preventing further expression of tau protein.

Herein, we report the development of a vectorized, species-specific artificial microRNA designed to target human tau transcripts. We demonstrate that amiRNA administration effectively reduces pathogenic tau in PS19 tauopathy mice at a pre-symptomatic stage, providing *in vivo* validation of a preclinical vector design. Subsequently, we bridge our preclinical vector findings with an intended clinical vector by evaluating efficacy at early disease onset. We further show that this therapeutic approach confers benefit even at advanced stages of disease, and we establish the durability of the intervention while determining a minimally effective dose in tauopathy mice. These results support the potential of vectorized amiRNA as a tau reduction strategy.

## RESULTS

### Generation of species-specific artificial miRNAs for tau transcription silencing

Commonly used transgenic mouse models of tauopathies show progressive accumulation of human tau while endogenous mouse tau remains independent of disease pathogenesis. Therefore, we hypothesized that efficacy assessments of reducing human pathogenic tau and not endogenous non-pathogenic mouse tau in a tauopathy mouse model requires a species-specific amiRNA. To generate species-specific amiRNAs capable of targeting all six tau isoforms expressed in the adult brain^28^, we performed sequence alignments of the 0N3R tau isoforms from human and mouse to identify regions of genetic divergence (Fig. 1a). Guide sequences were designed (11 human and 20 mouse targeted) and cloned into a murine U6 promoter (Pol III) driven expression plasmid containing a modified human miR-30a backbone. This enabled high-throughput screening of candidate amiRNAs by co-transfecting the miRNA expressing plasmid with a dual-luciferase plasmid encoding a 3’UTR tau cDNA tagged Renilla luciferase and a firefly luciferase as an internal transfection control (Fig. 1b). Primary screening (N = 2 independent experiments; n = 3 replicates/independent experiment) identified hit candidates from each species-specific library. The human amiRNA lead candidates, hTau5i, hTau6i, and hTau8i reduced human tau 3’UTR tagged Renilla expression >50% (58±2.2%; 66±1.9%; 62±4.2% respectively, Fig. 1c). Likewise, comparable targeting was observed in mouse amiRNA lead hit candidates, mTau2i, mTau7i, and mTau17i targeting mouse tau 3’UTR tagged Renilla expression (70±1.0%; 64±0.1%; 71±3.1% respectively, Fig 1d). Subsequently, secondary counter-screens (N = 2 independent experiments; n = 3 replicates/independent experiment) against the non-target species tau allowed us to confirm our lead hit amiRNA from each species-specific library (human: hTau5i, hTau6i, and hTau8i; mouse: mTau2i, mTau7i, and mTau17i) by eliminating cross-reactive sequences (Fig. 1c,d). This data supported further validation of our lead candidate amiRNAs.

**Figure 1.**
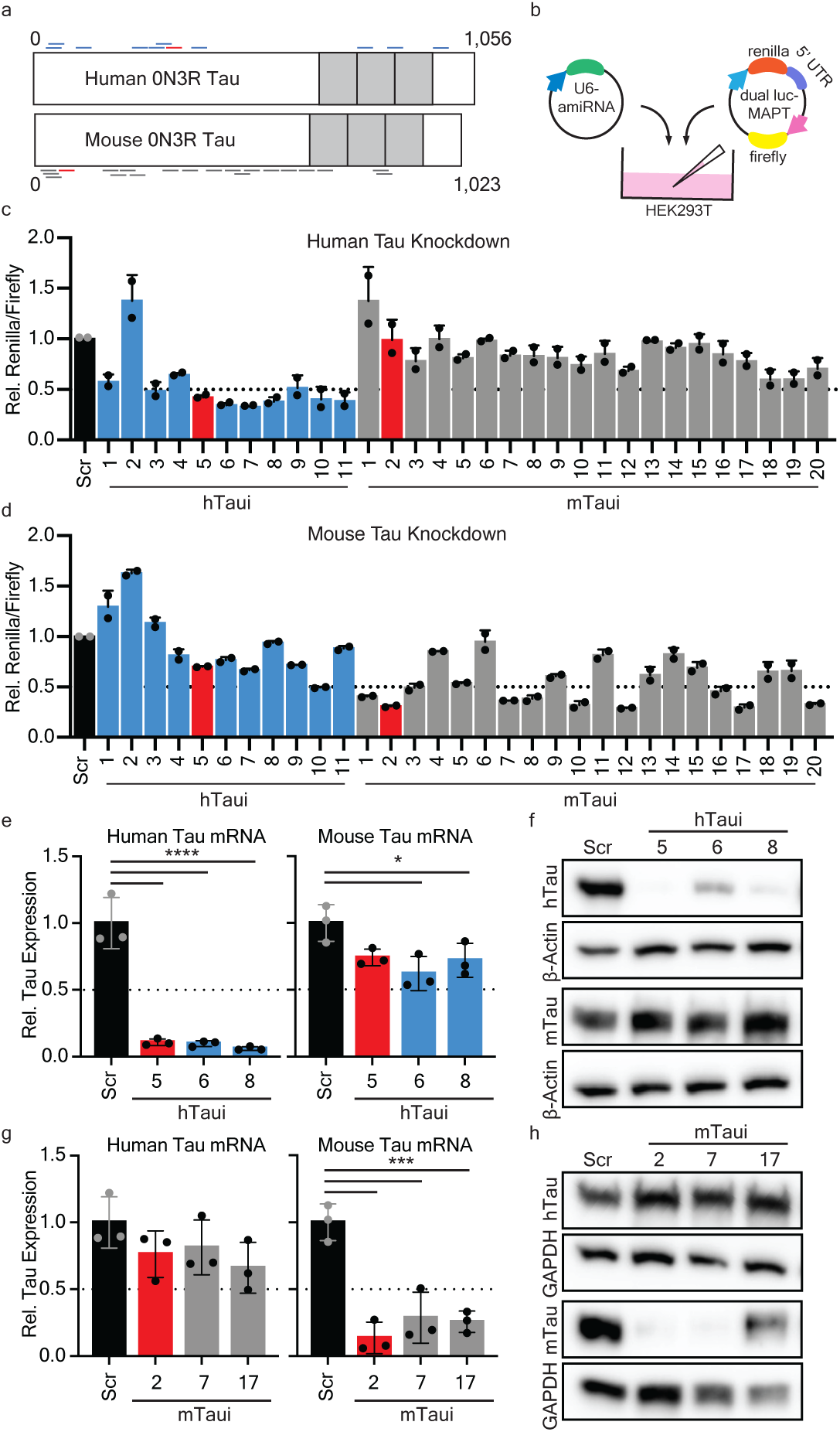
Screening rationally designed amiRNAs target and reduce microtubule associated protein tau in a species-specific manner. **a** Illustration of amiRNAs designed against human 0N3R tau (top - blue) and mouse 0N3R tau (bottom - gray). The lead candidate of each species-specific amiRNA is colored red. Gray boxes on the Tau gene represent the protein repeat domains. **b** Schematic representation of the amiRNAs screening paradigm using a dual luciferase reporter system. Plasmid containing a Firefly luciferase (internal transfection control) and Renilla containing 2N4R MAPT cDNA (“Human”) or 2N4R Mapt cDNA (“Mouse”) in the 3’ UTR of Renilla is co-transfected with a U6 driven amiRNA in HEK293T cells. **c,d** Histoplots of human tau **(c)** and mouse tau **(d)** silencing is shown relative to scramble miRNA control. Data points plotted represent the mean of N = 2 independent experiments; n = 3 replicates/independent experiment. **e-h** HEK293T cells were co-transfected with plasmids carrying the lead candidate species-specific tau amiRNAs and plasmids encoding either human tau (hTau) or mouse tau (mTau). Histoplots represent qPCR quantification of human (left) or mouse (right) tau mRNA when treated with human **(e)** or mouse **(g)** amiRNA lead candidates. Western blot images of lead human **(f)** or mouse **(h)** amiRNAs. n = 3 replicates. Data is displayed as Mean ±SD; One-way ANOVA with Tukey’s multiple comparisons; *p< 0.05, ***p< 0.001, ****p<0.0001. Dashed lines in histoplots represent the half maximal inhibitory value relative to the control group.

Lead amiRNA candidates were validated in cells that do not naturally produce tau protein, by co-expression with plasmids encoding human or mouse full-length tau (2N4R). Quantitative analysis of mRNA revealed that the top human tau-targeting amiRNA, hTau5i, achieved significant reduction of human tau transcripts by ∼90±2.4% (p<0.0001; one-way ANOVA; n=3/group) without affecting mouse tau mRNA levels (26±6.2%; p=0.0658; one-way ANOVA; n=3/group) (Fig. 1e, left). hTau6i and hTau8i reduced human tau comparable to hTau5i, however, statistically significant reduction in mouse tau mRNA was also observed (p<0.05; one-way ANOVA; n=3/group), indicating possible cross-reactive target engagement between species (Fig. 1e, right). Consistent with these findings, immunoblot analysis demonstrated robustly decreased human tau protein levels, while mouse tau levels remained unchanged (Fig. 1f). Similarly, the lead mouse tau-targeting amiRNA, mTau2i, selectively reduced mouse tau mRNA ∼87% (p<0.001; one-way ANOVA; n=3/group) and protein without impacting human tau (Fig. 1g,h), confirming the species specificity. Together, these functional and structural validations (Fig. 1 and Supplementary Fig. 1) establish a robust platform for precise, species-specific tau silencing.

### Validating in vivo silencing of human tau in a tauopathy mouse model

To support the advancement of our lead human tau-targeting amiRNA for *in vivo* evaluation we generated a preclinical vector encoding the lead amiRNA candidate hTau5i and eGFP (Fig. 2a). eGFP was included as a reporter protein to assess vector distribution in proof-of-concept studies. P301S mice are an established model of tauopathies that overexpresses human tau containing the disease-associated P301S mutation^29^. As early as 1-month of age, P301S mice demonstrate the capacity to template tau protein aggregation^7,30^. The preclinical vector was packaged into rAAV9 and administered intra-cisterna magna (ICM) to 3-mo-old wild-type (WT; vehicle n = 7; preclinical vector n = 8) and P301S tauopathy mice (vehicle n = 6; preclinical vector n = 7). Animals were monitored for body weight and survival over 3 mo, with no significant differences observed between vehicle- and vector-treated groups, indicating that our preclinical vector was well-tolerated in both genotypes (Fig. 2b,c). GFP immunostaining confirmed widespread vector distribution throughout the brain, with highest transduction near the injection site (brainstem and cerebellum) and decreasing levels of transduction in more distal regions (Fig. 2d).

**Figure 2.**
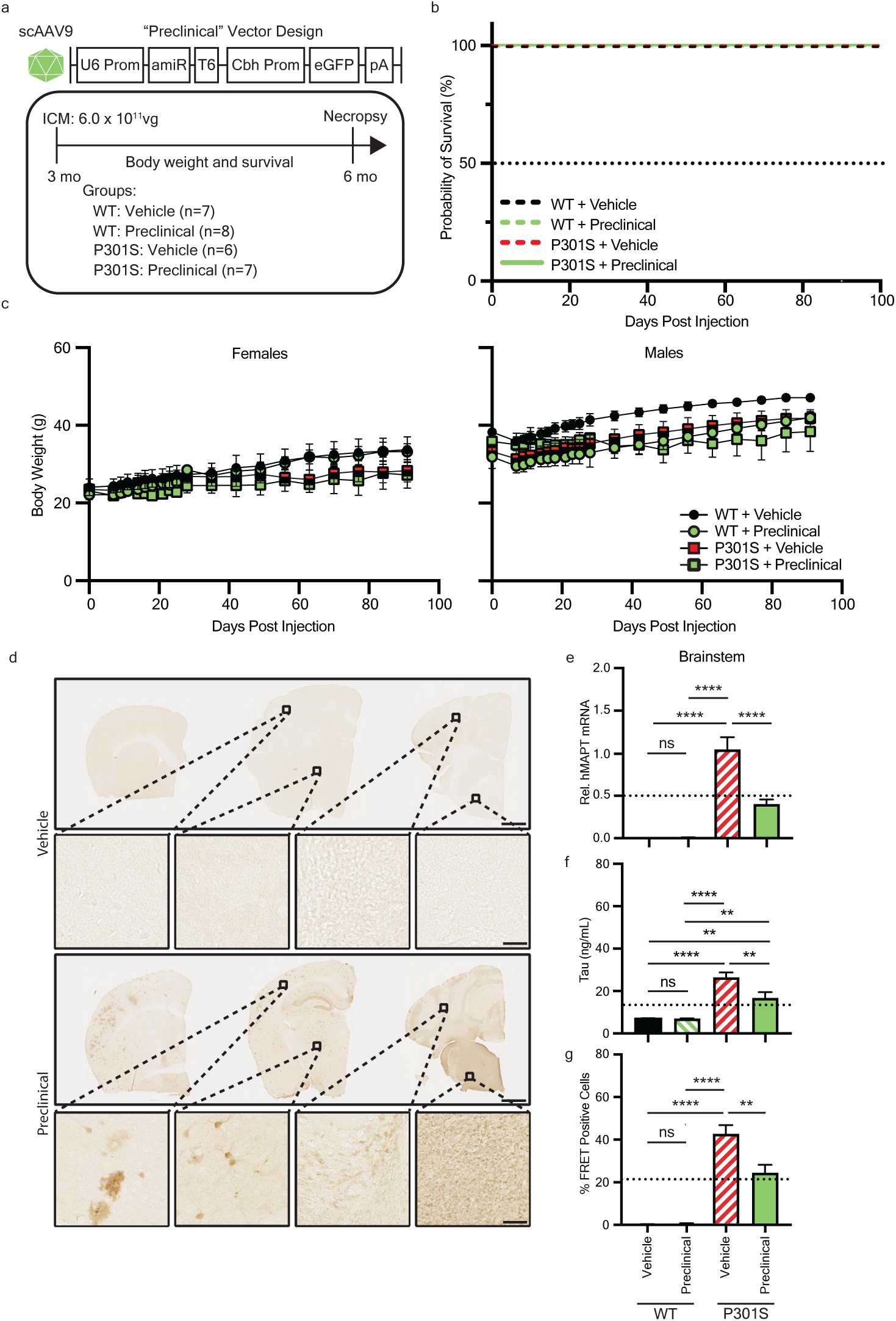
Pre-clinical vector administration in a tauopathy mouse model at a pre-symptomatic stage reduces human tau levels and seeding. **a** Graphical design of the “preclinical” human tau targeting amiRNA vector and experimental design. Three mo old WT or P301S mice were intra-cisterna magna (ICM) injected with pre-clinical vector (6.0 x 1011 vg) or vehicle control. **b,c** Mice were monitored weekly for survival **(b)** and weights **(c)** until study endpoint 3 mo post injection (6 mo of age). d Representative images of right coronal hemisphere of P301S brains stained for GFP in vehicle or pre-clinical vector injected animals. Scale bar = 1 mm (top) or 100 µm (middle and bottom). **e-g** Histoplots display measurement of tau mRNA **(e)**, protein **(f)**, and seeding activity **(g)** within brainstem samples. Data is displayed as Mean±SEM. One-way ANOVA with Tukey’s multiple comparisons. ns=not significant; **p< 0.01, ****p<0.0001. Dashed lines in histoplots represent the half maximal inhibitory value relative to the control group.

Quantitative PCR analysis demonstrated significant reduction of human tau mRNA in the brainstem of hTau5i treated animals compared to vehicle controls (p<0.0001; one-way ANOVA), confirming *in vivo* target engagement (Fig. 2e). Likewise, ELISA analysis revealed a corresponding decrease in total tau protein in P301S mice treated with hTau5i as compared to control treated P301S (p<0.01; one-way ANOVA). Endogenous mouse tau levels in WT animals remained unchanged, supporting the species specificity of hTau5i *in vivo* (Fig. 2f). Tau reduction was most pronounced in brain regions proximal to the injection site (brainstem and cerebellum), whereas more distal regions (e.g. cortex) exhibited less efficient knockdown, matching the observed pattern of vector expression via GFP staining (Fig. 2d and Supplementary Fig. 2). The presence of proteopathic tau correlates with formation of insoluble tau oligomers and aggregate-inducing seeds are present even at pre-symptomatic ages^7^. We applied mouse brain lysate to a tau biosensor cell line to measure proteopathic tau activity (“seeding”)^31^ and found a significant reduction in tau seeding in brainstem and cerebellum lysates (p<0.01; one-way ANOVA) from P301S mice treated with the preclinical vector as compared to vehicle treated (Fig. 2g and Supplementary Fig. 2b). Interestingly, we observed a small, trending decrease of seeding in the cortex (p=0.0563) although total tau protein levels were not affected (Supplementary Fig. 2).

These proof-of-concept studies validated the successful reduction of tau *in vivo* using our vectorized approach to deliver tau-reducing amiRNA therapies and supported further development and evaluation of a clinical version of our vector at later stages of disease.

### Bridging study of preclinical and clinical vectors at an early disease onset

To enable translation of our rAAV9/hTau5i approach for use in humans, we generated a “clinical” vector by removing the GFP reporter (Fig. 3a). Prior to rAAV9 packaging, we demonstrated that the preclinical and clinical vectors had comparable efficacy in targeting 4R and 3R tau in cells (Supplementary Fig. 3). To determine if there were adverse effects from overexpression of an amiRNA, we developed a non-targeting scrambled amiRNA to serve as a control. Sex-matched WT and P301S littermates were administered vehicle, scramble control, preclinical, or clinical vector at 6 mo of age, corresponding to early disease status (Fig. 3a). Mice were monitored for survival and body weight for 3 mo following injection. Body weight gain and survival were similar amongst all groups up to 9 mo of age, indicating that our scrambled and clinical vectors were well tolerated (Fig. 3b,c).

**Figure 3.**
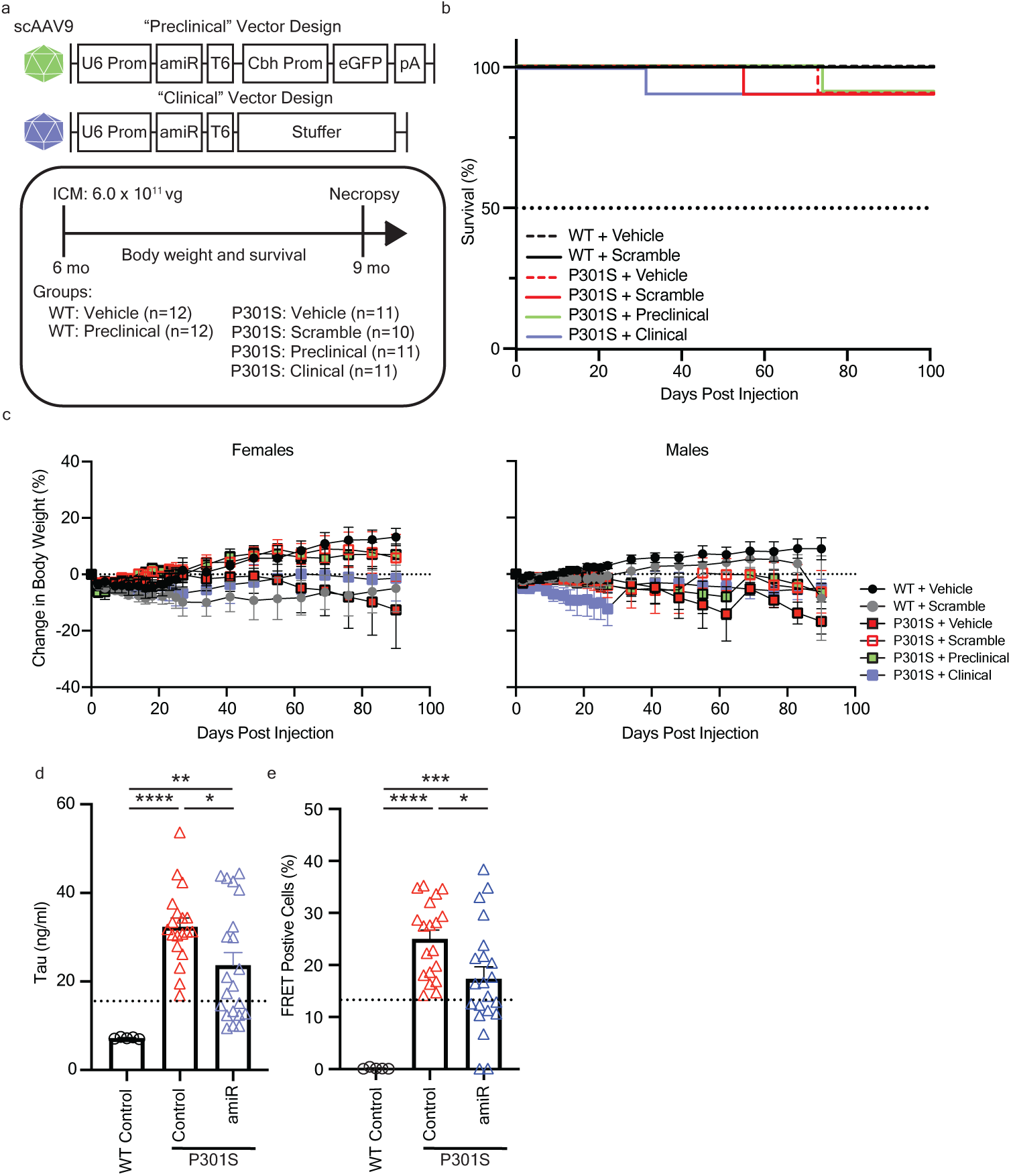
Bridging design of preclinical and clinical vectors shows comparable reduction of tau at an early disease intervention timepoint in tauopathy mice. **a** Graphical design of the “preclinical” (top) and “clinical” (bottom) human tau targeting miRNA (“hTau5i”) vectors. Experimental timeline following intra-cisterna magna (ICM) injection of 6 mo old WT and P301S mice. **b,c** In life assessments for survival **(b)** and percent body weight change **(c)** of mice for 3 mo post-injection. **d,e** Biochemical assessments at study endpoint (9 mo of age) for tau protein **(d)** and seeding activity **(e)** in brainstem samples. Data is displayed as Mean±SEM. One-way ANOVA with Tukey’s multiple comparisons; *p< 0.05, **p<0.01, ***p<0.001, ****p<0.0001. Dashed lines in histoplots represent the half maximal inhibitory value relative to the control group.

Analysis of tau protein showed no significant differences in tau reduction between vehicle and scramble controls or between preclinical and clinical vectors, therefore control groups and treatment groups were combined to increase statistical power. Consistent with results from intervention at 3 mo of age, both preclinical and clinical vector treatment at 6 mo of age significantly reduced human tau protein levels in the brainstem (p<0.05; one-way ANOVA; Fig. 3d) and cerebellum (p<0.001; one-way ANOVA; Supplementary Fig. 4a) of P301S mice as compared to control treated P301S mice when assessed 3 mo post-injection. Tau-targeting amiRNA treatment led to a significant decrease in tau seeding (p<0.05; one-way ANOVA; Fig. 3e) and cerebellum (p<0.01; one-way ANOVA; Supplementary Fig. 4b) samples. These bridging data support the therapeutic potential of our clinical vector design and indicate the potential of their efficacy when administered after the onset of disease.

### Late-stage intervention confers therapeutic benefit in a tauopathy mouse

Tauopathies are typically diagnosed after symptom onset and substantial neurodegeneration, therefore we interrogated whether our clinical amiRNA vector would provide therapeutic benefit at a late-stage disease timepoint by administering the amiRNA or scramble control vector to 9-mo-old P301S mice. Scramble control treated sex and age-matched WT littermates served as unaffected controls (Fig. 4a). In contrast to earlier assessments, P301S control mice exhibited significantly reduced survival and body weight compared to WT controls by 12 mo of age (Fig. 4b, c). Notably, clinical vector-treated P301S mice showed a two-fold improvement in survival relative to scramble controls (Fig. 4b). Sex dependent differences were observed with attenuated weight loss observed in male vector-treated P301S mice but not in females (Fig. 4c). To assess if prolonged survival was associated with improved quality of life, disease severity was determined using a disease score (see Methods). We found that hTau5i treatment improved quality of life, as indicated by a significantly higher proportion of mice with lower disease scores as compared to scramble treated mice (Fig. 4d). Late treatment was not sufficient to reverse disease progression as clinical amiRNA treated mice were significantly more affected than WT control mice (Fig. 4d).

**Figure 4.**
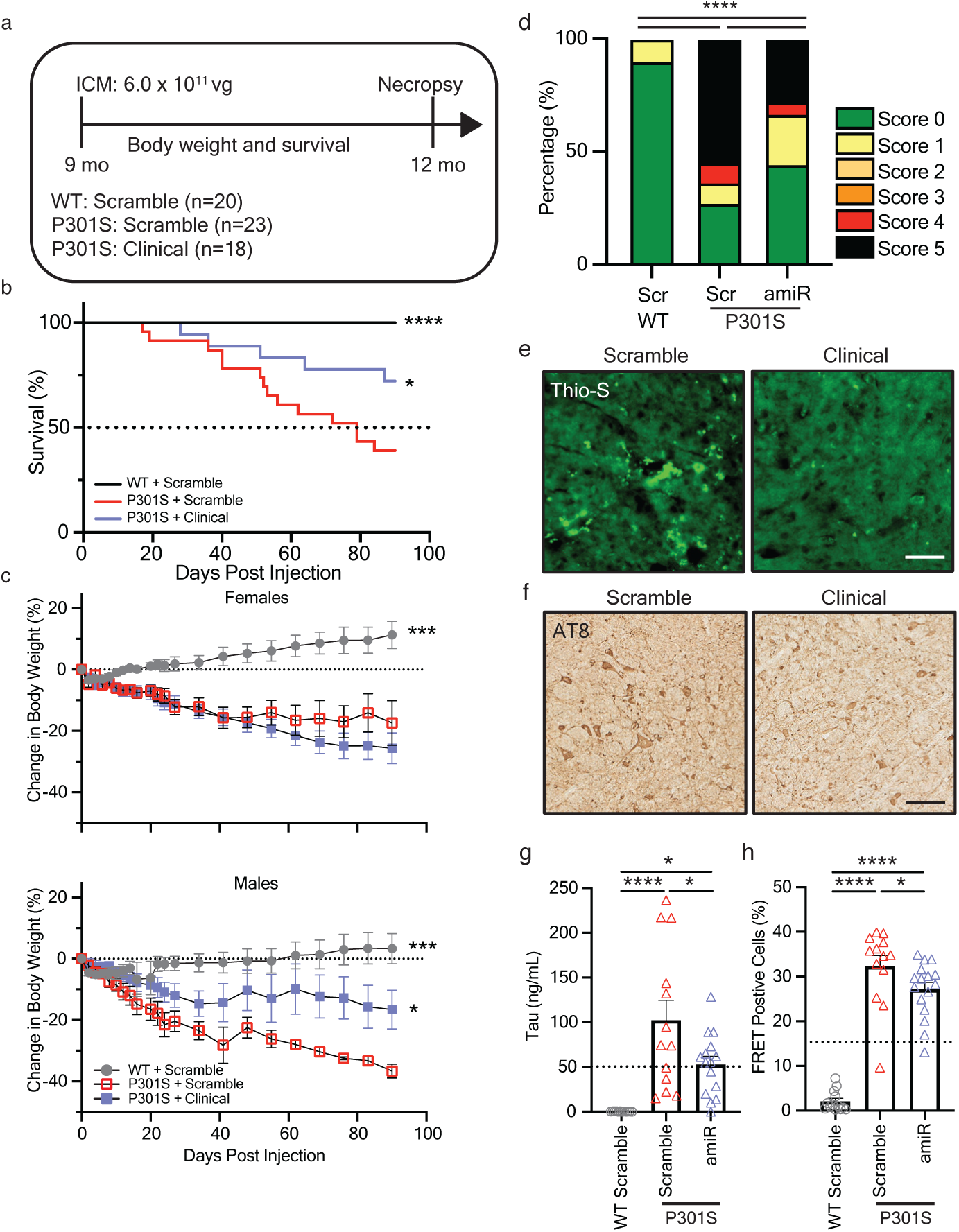
Late disease intervention with tau targeting amiRNA shows functional and pathological therapeutic benefit in tauopathy mice. **a** Experimental study design for testing treatment at late disease stage (9 mo of age). **b,c** Animals were monitored for survival **(b)** and percentage of weight change **(c)** until study endpoint 3 mo post injection (12 mo of age). **d** Quantification of disease score severity of animals injected with a scrambled control or tau targeting miRNA. **e,f** Representative images of Thio-S staining **(e)** in the brainstem, and AT8+ phospho-tau staining **(f)** within the cerebellum of scramble control injected or tau targeting amiRNA injected P301S littermates. Scale bar = 100 µm. **g,h** Quantification of tau protein **(g)** and seeding activity **(h)** in brainstem samples. Data is presented as Mean±SEM. One-way ANOVA with Tukey’s multiple comparisons for all panels except disease score **(d)** which was analyzed by Chi-square Goodness of Fit for observed vs. expected frequencies; *p< 0.05, ***p<0.0001. Dashed lines in histoplots represent the half maximal inhibitory value relative to the scramble control group.

In this older treatment cohort, we then determined if gene therapy treatment was effective at reversing pre-existing tau aggregates. Thioflavin S staining is a traditional method to identify accumulation of amyloid beta, tau, and other aggregating proteins in the brain. Likewise, post-mortem assessment of patients with cognitive impairment has categorized individuals with tauopathies by the degree of phosphorylated tau found through immunohistological staining of the brain using the AT8 antibody. Immunohistochemical analysis revealed qualitatively decreased Thioflavin S staining and qualitatively reduced AT8-positive tau pathology in the brains of vector-treated P301S mice compared to controls (Fig. 4e,f). Quantification of total tau protein by ELISA showed significant reduction in vector-treated animals (p<0.05; one-way ANOVA), accompanied by a corresponding decrease in pathogenic tau seeding activity (p<0.05; one-way ANOVA; Fig. 4g,h and Supplementary Fig. 5). Collectively, these findings demonstrate that late-stage intervention with a tau-targeting amiRNA confers therapeutic benefit in a tauopathy model.

### Sustained tau reduction at a minimally effective clinical vector dose

To evaluate the durability of therapeutic benefit and establish a minimally effective dose, P301S mice were treated at 6 mo of age with scramble control or three de-escalating doses (high, mid, low) of the clinical vector and monitored for 6 mo (Fig. 5a). Animals were randomly assigned to either an interim (3 mo post-injection) or endpoint (6 mo post-injection) group for analysis. Analysis of the interim group revealed that as expected, scramble treated P301S mice did not have significant early lethality or declines in body weight at 9 mo of age. Further, there were no significant differences between treatment groups in regards to survival and change in body weight (Fig. 5b-d), in agreement with our prior study (Fig. 3b-d). Analysis of the endpoint group revealed that only the high dose conferred significant survival benefit relative to controls (Fig. 5e), and recapitulated the survival advantage observed in our prior 3 mo duration study (Fig. 4b). Additionally, the high dose attenuated body weight decline in male mice in the endpoint group (p = 0.075; two-way ANOVA Fig. 5f). In this study group, scrambled P301S mice did not decline in body weight, which was similar in the hTau5i treatment groups (Fig. 5g).

**Figure 5.**
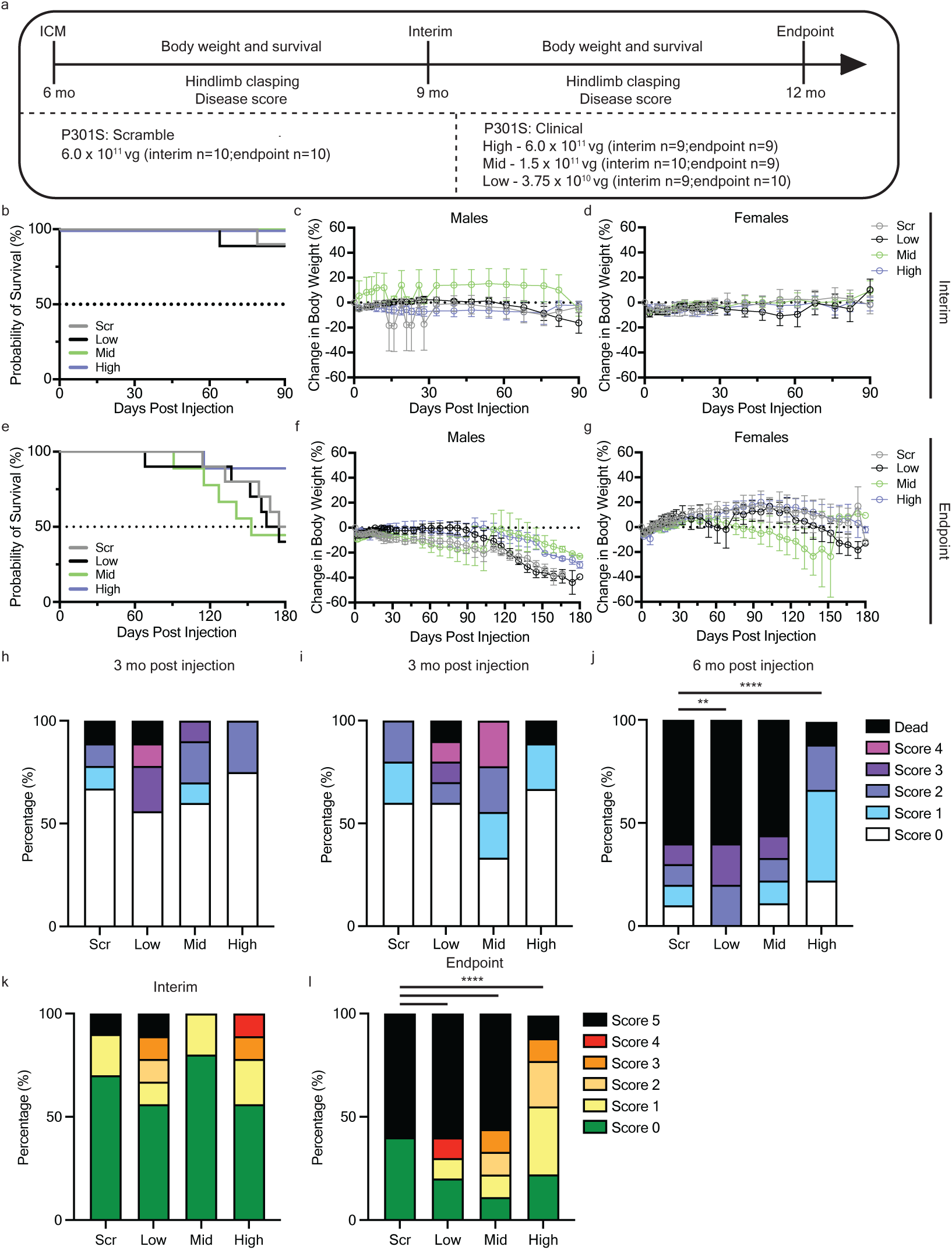
Dosing studies reveal a minimal effective dose and treatment durability. **a** Experimental design for dosing 6 mo old P301S mice for interim (3 mo post-injection) and endpoint (6 mo post-injection) analysis groups. **b-g** Interim **(b-d)** and endpoint **(e-g)** animals were monitored for survival **(b,e)** and body weight **(c,d,f,g)**. **h-j** Mice were assessed for hindlimb clasping **(d)** 3 mo post injection for interim **(h)** and endpoint **(i)** groups, and 6 mo post injection for endpoint animals **(j)**. **k,l** Interim **(k)** and endpoint **(l)** animals were evaluated for disease severity prior to study endpoint. Disease score and hindlimb clasping were analyzed by Chi-square Goodness of Fit for observed vs. expected frequencies; **p< 0.01, ****p<0.0001.

Hindlimb clasping correlates with disease progression in neurodegenerative mouse models, such as Alzheimer’s disease, Parkinson’s disease and cerebellar ataxias^32^. In P301S tauopathy mice lower limb paresis begins to develop at approximately 7-9 mo of age, which progresses towards early mortality. Analysis of hindlimb clasping in the interim group at 3 mo post injection showed that ∼44% of low dose, ∼10% of mid dose, and 0% of high dose treated P301S mice had moderate to severe disease scores (scores 3-4) or early death as compared to 10% of scrambled treated P301S mice (Fig. 5h). Assessment of endpoint group animals at 3 mo post injection displayed relatively similar trends to the interim cohort (Fig. 5i). Analysis of hindlimb clasping 6 mo post injection (12 mo of age) revealed that only the high-dose resulted in sustained functional improvement as 89% of high dose treated mice had none to mild-moderate hindlimb clasping (scores 0-2) as compared to scramble (30%), low (20%), and mid (33%) dose treated mice (Fig. 5j). Moreover, low dose treated P301S mice were more severely affected than scramble treated mice with all surviving mice being moderately to severely affected.

Analysis of disease severity in the interim group was not indicative of therapeutic benefit, as at this age, score-driving criteria were minimal in all treatment groups (Fig. 5k). In contrast, disease severity scores at endpoint culminated in 60% of scramble treated P301S mice with severe to end-stage disease severity (scores 4-5) (Fig. 5l). Treatment with either low or mid dose tau amiRNA did not improve disease progression or severity as surviving mice were significantly more affected compared to the scramble treated group. In contrast, treatment with the high dose at 6 months of age significantly decreased the amount of severe and end-stage affected P301S mice to 11% (Fig. 5e) which provided improved benefit as compared to P301S mice treated at 9 months of age (Fig. 4d). Together, in-life assessments support the high dose as the minimally effective therapeutic dose in P301S mice.

We then assessed tau protein in brain tissues from interim and endpoint animals. The high dose showed a sustained reduction of total tau levels within the brainstem and cerebellum to ∼45 and ∼59 ng/mL in the interim and endpoint groups, respectively, as compared to the scramble treatment, which were ∼59 and ∼93 ng/mL, respectively (Supplementary Fig. 6). Tau protein levels were highly variable in these cohorts, however, so we did not observe a significant dose-dependent effect in treated mice. Assessment of tau seeding in brain tissue showed that in the interim group there was a significant reduction in seeding activity in low, mid, and high tau amiRNA dose groups (Supplementary Fig. 6), At endpoint, seeding in high dose treated mice was reduced to ∼65% of scramble treated mouse seeding levels, although this reduction was not significant (Supplementary Fig. 6). In contrast, low and mid dose treated P301S mice had similar seeding levels as the scrambled treated mice (Supplementary Fig. 6). These findings indicate that in P301S mice, sustained tau reduction and therapeutic benefit were achieved with only the highest tested clinical vector dose.

## DISCUSSION

Currently, there are no approved disease-modifying therapies for tauopathies, and available treatments are limited to symptomatic relief^33^. Previous studies have demonstrated the therapeutic potential of reducing tau protein levels using antisense oligonucleotides (ASOs)^34^. While promising, this therapeutic modality requires repeat administration, due to their relatively short half-life and transient effects. The practical implementation of repetitive ASO therapy procedures poses additional burdens to patients with age-related neurodegenerative diseases. Our study aimed to address these limitations by vectorizing an amiRNA targeting human tau and delivering it via a single dose of rAAV9 for long-lasting therapeutic benefit. We demonstrated that our rAAV9/hTau5i approach achieved significant and durable reduction of tau protein across pre-, early, and late symptomatic intervention timepoints in a P301S tauopathy mouse model. Notably, therapeutic benefit was observed regardless of age of administration. Further, tau reduction correlated with decreased aggregation activity, improved probability of survival, and enhanced quality of life as assessed by hindlimb clasping and disease severity scores. These findings provide proof-of-concept that a single administration of a vectorized tau-reducing agent can confer sustained disease modification, even when delivered after the onset of neurodegeneration.

The P301S mouse model is widely used in the tauopathy field, although limitations of this model include overexpressed tau protein levels that far exceeding physiological expression^29^, and there is significant variability in tau pathogenesis and progression between litters and different colonies^7,18,35^. To mitigate these issues, we randomized treatment assignments within littermates, employed large cohorts, and assessed multiple treatment groups to confirm the reproducibility and robustness of the drug effect. The reversal of tau pathology that we achieved with rAAV9/hTau5i was consistent with previous studies using ASOs to reduce human tau in the same mouse model and with similar age of treatments^18^. While DeVos et al. reported a similar reduction of tau seeding in groups of mice treated at 6 or 9 mo of age and assessed 3 mo later^18^, we found a greater reduction of tau seeding with treatment at 6 mo of age (Fig. 3e and Supplemental Fig. 4b) as compared to treatment at 9 mo of age (Fig. 4h and Supplementary Fig. 5b). As there are more tau aggregates present at the time of treatment in 9 mo old mice, this is consistent with expectations and differences noted in the DeVos study may have been a result of biological variability, as they suggested. We found vectorized tau reduction prevented functional decline and improved quality-of-life as assessed by hind-limb clasping, disease scoring, and premature death. Further, improvements in quality of life were greater when P301S mice were treated at 6 months of age as compared to treatment at 9 months of age. However, a limitation of this current work is that changes in cognition were not assessed with treatment and future work will aim to address this shortcoming. Additional studies utilizing relevant models expressing endogenous tau levels may also be more informative on dosing considerations.

Our vector distribution results were consistent with other studies^36–38^ where intra-cisterna magna delivery of AAV9 in adult mice resulted in vector transduction highest in areas closest to the injection site, such as the brainstem and cerebellum. As expected, areas with the highest amount of vector transduction, such as the brainstem and cerebellum, showed greater reduction of tau protein levels as compared to more distal regions, such as the midbrain, which had lower levels of vector transduction. In these regions, the level of tau protein reduction correlated with a reduction in tau seeding. Interestingly though, tau protein levels were unchanged in the cortex, yet tau seeding was reduced to a small extent. As tau seeds are thought to move between cells and synapses in the brain^5,7,8^, reduction of tau in the hindbrain may have slowed the spread of pathological forms of tau along networks to distal regions, such as the cortex. This suggests that reduction of tau in one area of the brain may have cell non-autonomous benefits for connected regions. Additional studies are needed to test this.

Our study demonstrated that rAAV9/hTau5i was well tolerated in both WT and tauopathy mice as assessed by body weight, gross behavior, and survival. It should be noted that in P301S mice, while human tau was significantly reduced, endogenous mouse tau levels were unchanged. Prior work utilizing a vectorized short hairpin RNA targeting endogenous mouse tau demonstrated learning and memory deficits when mouse tau was acutely reduced in the hippocampus^39^. Additionally, assessments of tau knockout reported increased anxiety and impaired contextual and cued fear memory^40^. Further investigation of safety of our vectorized amiRNA tau reduction approach should be performed either within a more physiologically relevant mouse model, such as human tau knock-in mice^41,42^, or by using our mouse tau targeting amiRNA in WT mice. However, a study utilizing tau targeting antisense oligonucleotides reported similar reduction in tau within P301S mice (∼50%), similar to our findings, and demonstrated reversal of tau pathology^18^. Moreover, RNA silencing machinery inherently has been evolutionarily pressured to have a certain degree of promiscuity^22,43–46^. Future development of this lead artificial miRNA candidate will warrant evaluation of off-target gene silencing.

A prohibitive barrier in the application of CNS gene therapies remains wide-spread and efficient delivery of therapeutic agents to critically affected brain regions^20^. Spatiotemporal distribution of tau lesions across tauopathies have diverse heterogeneity^47^, in that cisterna magna delivery of an AAV-mediated tau modifying treatment may not be efficient for all disorders. Although rAAV9 remains the dominant vector due to its favorable safety profile, recent advances in engineered AAV capsids^48,49^ and emerging delivery technologies, such as MRI-guided focused ultrasound (FUS)^50–52^, offer promising avenues to enhance CNS transduction, reduce peripheral biodistribution, enable less invasive delivery, and lower the required vector dose to achieve therapeutic benefit. Future work integrating next-generation capsids and FUS could further improve the therapeutic index of our approach and enable more uniform targeting of affected regions in diverse tauopathies.

It is important to consider that the age at which AAV9 is administered may impact the degree of CNS transduction achieved. Previous work from our group demonstrated that intrathecal delivery of AAV9 in mice results in age-dependent differences in transduction efficiency, with younger animals showing more widespread vector distribution^53^. However, that prior study did not assess intra-cisterna magna delivery or older ages, such as 6 or 9 mo, as performed in the present study. Therefore, it remains possible that advanced age could present additional challenges for effective CNS transduction via intra-cisterna magna administration. Further investigations are required to systematically evaluate the impact of age on vector biodistribution and therapeutic efficacy following intra-cisterna magna delivery, particularly in the context of late-stage intervention for tauopathies.

In summary, our study provides a framework for the development of translational, vectorized amiRNA therapies for tauopathies and potentially other neurodegenerative diseases. By demonstrating efficacy at both early and late disease stages, establishing a minimally effective dose, and validating the durability of benefit, our findings support further development toward clinical translation. Collectively, these results advance the field toward a disease-modifying therapy for tauopathies, with the potential for broad impact on age-related neurodegenerative disorders.

## METHODS

### Sequence generation and cloning

Designs for amiRNAs targeting microtubule associated protein tau were generated using the Harper shuttle predictor v1.0 as previously described^26^. In brief, 0N3R isoform of mouse *Mapt* or human MAPT coding sequence was used as input into the Harper shuttle predictor. Output sequences were assessed for species specificity and targeting of all 6 isoforms expressed in the CNS using NCBI BLAST. Hairpin secondary structures and cut sites of lead human and mouse amiRNAs were predicted using the Unified Nucleic Acid Folding (UNAFold) software package^54^. The amiRNA sequences were cloned into a U6-miR30 expression cassette as previously described^25^.

### Luciferase Assay

amiRNAs were screened using a Dual Luciferase Reporter Assay (Promega). The dual luciferase reporter plasmid was modified from PsiCheck2 (Promega) containing a firefly luciferase, serving as an internal transfection control, and the mouse or human 2N4R MAPT gene cloned downstream of the Renilla luciferase stop codon serving as a 3’ UTR. HEK293T cells were co-transfected using Lipofectamine 2000 (Invitrogen) and the dual luciferase reporter plasmid and individual MAPT targeting amiRNA plasmids in a 1:5 molar ratio (target:amiRNA). Screens were conducted as 2 independent experiments and triplicate data points were averaged per individual experiment. Results of the luciferase screens are reported as a ratio of Renilla to Firefly luciferase activity ±SD for all combined experiments.

### Immunoblotting

Total protein was extracted from HEK293T cells transiently co-transfected with a CMV.MAPT vector (mouse or human) and candidate amiRNA using lysis buffer containing 50 mM Tris-HCl (pH 7.5), 274 mM NaCl, 5 mM KCl, 5 mM EDTA, 1% Triton-X100, 1X protease inhibitor (Roche), and 1X phosphatase inhibitor cocktail (Sigma). Protein concentration was determined by Pierce BCA assay (Invitrogen). 10 μg of total protein per sample were separated on 10% SDS-PAGE gels (Biorad). Protein was transferred onto PVDF membranes using a turbo-blot transfer system (Biorad). Blots were blocked with 5% milk in TBS-T for 1 h at RT then incubated overnight at 4 °C in 5% milk-TBST containing primary antibodies targeting either mouse monoclonal TAU-5 antibody (1:1000, ThermoScientific, AHB0042), rabbit monoclonal GAPDH (1:1000, Meridian, H86504M), or rabbit monoclonal β-actin (1:1000, Cell Signaling, 4970S) followed by HRP-conjugated goat anti-mouse or goat anti-rabbit secondary antibodies (1:10,000, Jackson Immuno, 115-035-146 or 111-035-144) at RT for 1 h. Blots were developed using enhanced chemiluminescence (ThermoScientific) on a G:Box Chemi XX6 system using GeneSys software (Syngene). β-actin or GAPDH were used as housekeeping loading controls.

### Experimental Animals

All procedures were performed in accordance with protocols approved by the Institutional Animal Care and Use Committee of the University of Texas Southwestern Medical Center (UT Southwestern), an AAALAC-accredited facility. All animal protocols were conformed to the NIH Guide for the Care and Use of Laboratory Animals. A colony of PS19 mice were maintained by breeding male mice carrying the 1N4R human tau isoform with the P301S mutation with female B6C3F1/J mice (Jax stock: 100010). Sex and age matched non-transgenic littermates served as WT controls where indicated. Mouse genotypes were determined by PCR on tail genomic DNA using the following primer pairs. (P301S-F: GGGGACACGTCTCCACGGCATCTCAGCAATGTCTCC; P301S-R:GCCTAGACCACGAGAATGCGAAGGAACAAG; Fabpi5’: TGGACAGGACTGGACCTCTGCTTTCCTAGA; Fabpi3’: TAGAGCTTTGCCACATCACAGGTCATTCAG)

### Intracisterna Magna Injection

3-, 6-, or 9-mo-old mice were randomized, and treatment groups were sex matched within littermates. Intra-cisterna magna injections were carried out as previously described^38^. Briefly, surgical procedures were conducted using aseptic technique under anesthesia induced and maintained by 2% vaporized isoflurane. Following hair removal, a small incision was made in the back of the scalp exposing the top of the head and neck muscle. Lidocaine was used as a topical anesthetic and carprofen (5 mg/kg) was administered via subcutaneous injection as an analgesic at the time of surgery and up to 48 h after. A 50 μL Hamilton syringe was loaded with 10 μL of virus and used for freehand single bolus injection into the cisterna magna space. The incision was sealed using 3M Vetbond surgical adhesive and mice were monitored within a recovery chamber until ambulatory. Mice were monitored in their home cages for 48 h post-surgery.

### Hindlimb Clasping and Disease Scoring

Mice were evaluated and scored for the degree of hindlimb clasping and disease progression as indicators of tauopathy severity. Hindlimb clasping was evaluated on an increasing scale of 0 to 5 where 0 is unaffected with complete splay of hindlimbs when grasped by the tail. A score of 1 indicates one or both hindlimbs exhibit mild, partial retraction towards the body. A score of 2 is mild-moderate where both hindlimbs are partially retracted 50-75% towards the midline of the body. 3 is moderate where neither hindlimb extends beyond the perimeter of the body. 4 is moderate-severe where neither hindlimb extends more than 10% from the midline of the body. A score of 5 is severe with full hindlimb clasping toward the midline and no evidence of extension.

Disease condition severity was determined for each mouse using a “disease score”. Mice with no discernible symptoms were scored as 0. Mice beginning to exhibit a hunched back were scored as 1. Mice which progressed to tremors and mild weight loss (less than 10%) were scored as 2. Mice exhibiting early paresis with moderate weight loss (10-15%) but still mobile were scored as 3. Mice whose symptoms progressed to partial paralysis with corresponding weight loss over 20% were scored as 4. Mice found dead or in conditions necessitating early humane euthanasia (e.g. full paralysis, inability to upright, etc.) were scored as 5.

### Tissue collection

Mice were deeply anesthetized with avertin and perfused with ice-cold PBS. Brains were rapidly dissected and divided into hemispheres. One hemisphere was sub-dissected into the following regions: brainstem, cerebellum, hippocampus, midbrain, cortex. Each region was rapidly homogenized and separated into 2 tubes designated for either protein or RNA analysis. Tubes were rapidly frozen on dry-ice and stored at −80 °C. The opposing hemisphere was drop-fixed in 4% paraformaldehyde for two days at RT, then prepared for cryopreservation by successive rounds of 10%, 20%, and 30% sucrose for 24 h each and stored at −80 °C for subsequent histological analysis.

### Quantitative RT-PCR anlaysis

Total RNA was purified using Qiazol (Qiagen) for mouse tissues or RNeasy plus mini kit (Qiagen) for cell samples and cDNA was synthesized using 1 µg of total DNase I-treated RNA using random hexamer primers provided in Transcriptor First Strand cDNA synthesis kit (Roche) per manufacturer’s specifications. Measurements for human or mouse MAPT were quantified using PowerUp SYBR Green Master Mix (Applied Biosciences) with a StepOne Plus (Applied Biosciences) thermocyler per manufacturer’s specification.

### Protein Analysis

To measure tau protein expression and seeding activity, brain tissues were prepared as follows. Brain subregions were each homogenized in 10 µl lysis buffer per mg of tissue weight (50 mM Tris-HCl [pH 8.0], 274 mM NaCl, 5 mM KCl, supplemented with protease inhibitor [Complete, Roche] and 1% phosphatase inhibitor cocktail [Sigma]) using a TissueLyser LT Adapter (Qiagen) at 40 Hz for 3 min. Equal volumes of homogenate were transferred to Beckman-Coulter centrifuge tubes and centrifuged at 150,000 x g on an Optima MAX-XP Ultracentrifuge (Beckman-Coulter) for 15 min at 4 °C. Supernatant was collected, and protein concentration was measured using Pierce Bicinchoninic Acid (BCA) Protein Assay Kit (Thermo Scientific).

### Tau ELISA

Tau protein levels were measured on a Tau5-HT7 sandwich enzyme-linked immunosorbent assay (ELISA) performed as previously described^55^ with minor modifications. 96–half area–well high binding plates (Corning) were coated with the TAU-5 antibody (AHB0042, LIfe Technologies) diluted in sodium bicarbonate buffer for 48 h at 4°C, followed by decanting and blocking with SuperBlock™ Blocking Buffer (ThermoFisher) for 2 h at 37°C. Recombinant 1N4R human tau (Boston Biochem, Cat. # SP-501) was diluted in 20% SuperBlock-TBS to generate a standard curve. 50 μL of 1.5 μg total protein diluted in 20% SuperBlock-TB or standards were added to each well and incubated overnight at 4°C. The next day, samples and standards were decanted and washed in 1X PBS followed by incubation in biotinylated HT7 (diluted 1:3000 in 20% Superblock-TBS, MN1000B, Thermo Scientific) for 2 h followed by incubation in streptavidin poly–horseradish peroxidase–40 (Fitzgerald). Super Slow ELISA TMB (Sigma-Aldrich) was added to wells for 30 min followed by ELISA Stop Solution (Bethyl). Absorbance at 450nm was read on an Epoch Microplate Spectrophotometer (BioTek).

### Tau Seeding Assay

In vitro tau seeding activity was carried out and measured as previously described by Holmes et al 2014^7,56^. Briefly, HEK-293T cells which stably express the repeat domain of tau fused with CFP or YFP were plated in 96-well plates at a density of 2.5 x 10^4^ cells in 180 µL culture media per well. 24 h later, at roughly 60% confluence, cells were seeded with brain lysates (30 µg per well). The transfection complexes were generated by incubating Opti–Minimum Essential Medium (Gibco) with the lysate and Lipofectamine 2000 (Invitrogen) for a final volume of 10 µL per well for 20 min at room temperature. 48 h later, cells were harvested with 0.25% trypsin followed by fixation in 4% paraformaldehyde (Electron Microscopy Sciences) for 10 min at RT. Fixed cells were resuspended in flow cytometry buffer (HBSS plus 1% FBS and 1 mM EDTA). To quantify tau seeding, FRET flow cytometry was performed using LSR Fortessa flow cytometer (BD Bioscience). To measure the mCerulean3 and mClover3 signal, laser excitation of cells was performed at 405 nm and 488 nm, respectively, followed by capture of fluorescence using 405/50 nm and 525/50 nm filters, respectively. FRET signal was measured using excitation at 405 nm followed by fluorescent capture using a 525/50 nm filter. FRET signal was quantified using a gating strategy where the mCerulean3 bleed-through into the mClover3 and FRET channels analyzed using FlowJo analysis software (version10.10.0, Beckman Dickson Co.), as previously described^56^. FRET measures are reported as the percentage of FRET-positive cells in all analyses.

### AT8 staining

Fixed hemispheres were mounted on a Leica Frozen Microtome (VT 1000S; Leica Biosystems Inc., Buffalo Grove, IL) and serially sliced into 30 µm brain sections. Serial sections were mounted onto slides and dried overnight. Slides were blocked with normal goat serum (101098-382, VWR) at room temperature for 1 h then incubated in primary AT8 antibody diluted in PBS (1:1000, MN1020B, Invitrogen) at 4 °C overnight. Slides were maintained under low light conditions for incubation in secondary antibody diluted in PBS (1:500 α-rabbit 488 & 1:500 Strepavidin-594) at RT for 1 h followed by incubation with DAPI (Sigma-Aldrich) for 10 sec. Sections were cover-slipped with aqua-polymount and dried before imaging at 20X on a Hamamatsu NanoZoomer 2.0 HT. All sections were stained on the same day and imaged together to prevent batch variability. The percent area identified as AT8–positive Tau was calculated using the Area Quantification algorithm in Halo Image Analysis software (v. 3.4.2986.1780, Indica Labs, Albuquerque, NM).

### Thioflavin S staining

Serial 30 µm brain sections were mounted onto slides and dried overnight. Slides were incubated with 0.05% Thioflavin S (Sigma-Aldrich) in 50% ethanol for 8 min in the dark, dipped into 80% ethanol for 15 sec, and then cover-slipped with aqua-polymount mounting medium with propidium iodide. Slides were scanned at 20x by light-microscopy using a Zeiss Axioscan Z1. All sections were stained on the same day and imaged together to avoid batch variability.

### Statistical Analysis

All experiments were analyzed and graphed using Graphpad Prism (version 10.5, GraphPad Software). Differences between two groups were determined by unpaired t-test. Differences between the means of multiple groups were determined via One-Way ANOVA with Tukey post-hoc multiple comparisons. Survival was assessed by the Log-rank (Mantel Cox) test. Differences in the change in body weight between treatment groups were evaluated by 2-way ANOVA Mixed Effects Restricted Maximum Likelihood Model followed by Tukey’s multiple comparisons. The Chi-square Goodness of Fit test for differences between observed and expected values was applied to disease severity and hindlimb clasping scores. For purposes of analysis, scores 1-2 were categorized as mild-moderate and scores 3-4 were categorized as moderate-severe. Statistical significance was set at p < 0.05.

## ACKNOWLEDGEMENTS

We would like to thank Marc Diamond, M.D. for PS19 breeder mice and HEK293T tau biosensor cells. We thank the University of Texas Southwestern Medical Center’s Whole Brain Microscopy Facility, RRID:SCR_017949. We thank Scott Harper, Ph. D. for the Harper shuttle predictor and U6T6 expression plasmid. This work was supported by the Crowley Foundation (RMB), Broughton Foundation (RMB), the National Institutes of Health 1R01AG078417 (RMB), Taysha Gene Therapies (RMB), the Peter O’Donnell Jr. Brain Institute’s Neural Scientist Training Program (ITG), and Mechanisms of Disease & Translational Science Training NIH T32 Grant GM109776 (ITG).

## AUTHOR CONTRIBUTIONS

RMB designed and conceptualized the study. ITG, SH, KK, and KP performed experiments. ITG, BS, and RMB analyzed data. RMB supervised the study. ITG, BS, and RMB wrote the manuscript. All authors reviewed the manuscript.

## DATA AVAILABILITY

Data supporting the findings of this manuscript are available from the corresponding author upon request. The dataset generated in this study, which is necessary to interpret, verify and extend the research in the article, is provided in the Supplementary Information/Source Data file. Source data are provided with this paper.

---

**Supplementary Figure 1.**
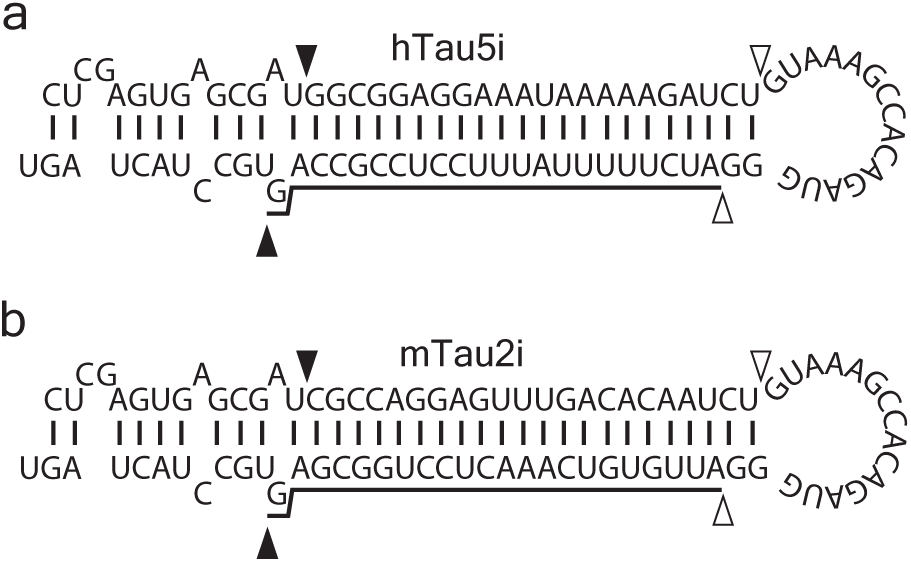
In silico prediction of artificial tau miRNAs. **a,b** UNAFOLD hairpin 2 dimensional structures and sequences of lead human **(a)** and mouse **(b)** artificial miRNAs. Black and gray arrowheads indicate Drosha and Dicer cut sites respectively in the modified human miR-30a cassette and the underlined area indicates the guide miRNA strand sequence.

**Supplementary Figure 2.**
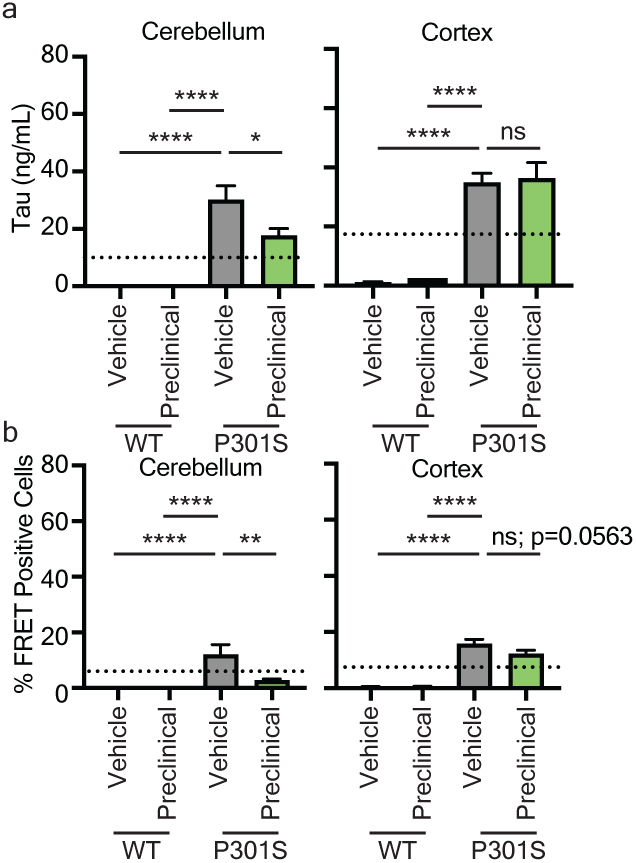
3-month efficacy data. **a** ELISA quantification of 1N4R human P301S tau cerebellum and cortex; **b** Soluble tau seeding of cerebellum and cortex lysates from wildtype and P301S littermates using HEK293T tau biosensors. Mean±SD; One-way ANOVA with Tukey’s multiple comparisons; * p< 0.05; ** p< 0.01; ****p<0.0001. Dashed lines in histoplots represent the half maximal inhibitory value relative to the control group.

**Supplementary Figure 3.**
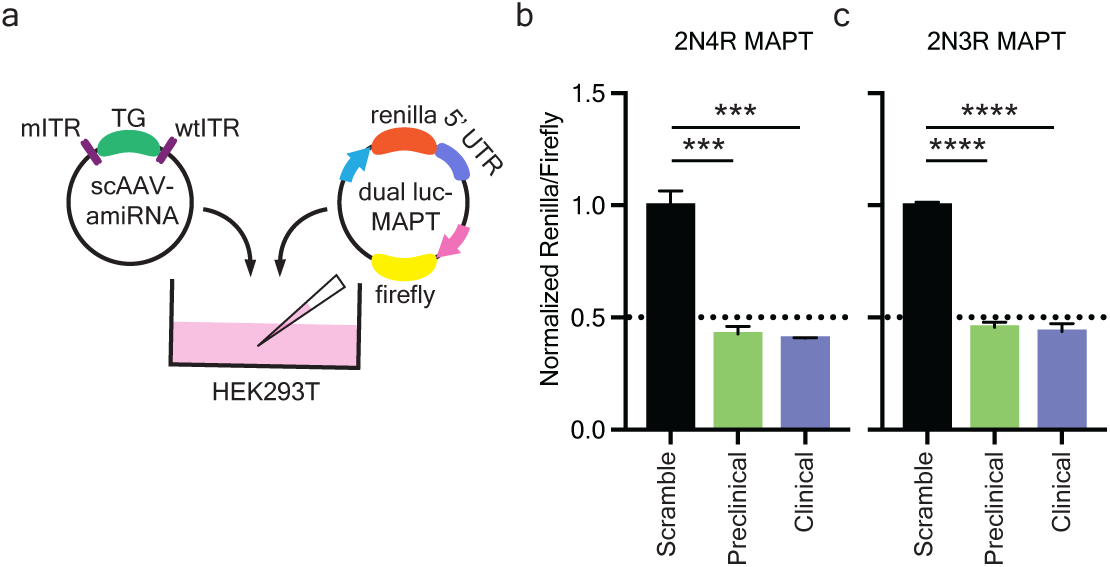
Preclinical and clinical vector designs display comparable total tau gene reduction. **a** Illustration of experimental paradigm of the self-complimentary AAV constructs co-transfected into microwell plates with the MAPT dual luciferase screening constructs; **b,c** Histoplots illustrate the reduction of human 2N4R **(b)** and 2N3R **(c)** MAPT isoform knockdown using a dual luciferase reporter system. Mean±SD; One-way ANOVA with Tukey’s multiple comparisons; *** p< 0.001; ****p<0.0001. Dashed lines in histoplots represent the half maximal inhibitory value relative to the control group.

**Supplementary Figure 4.**
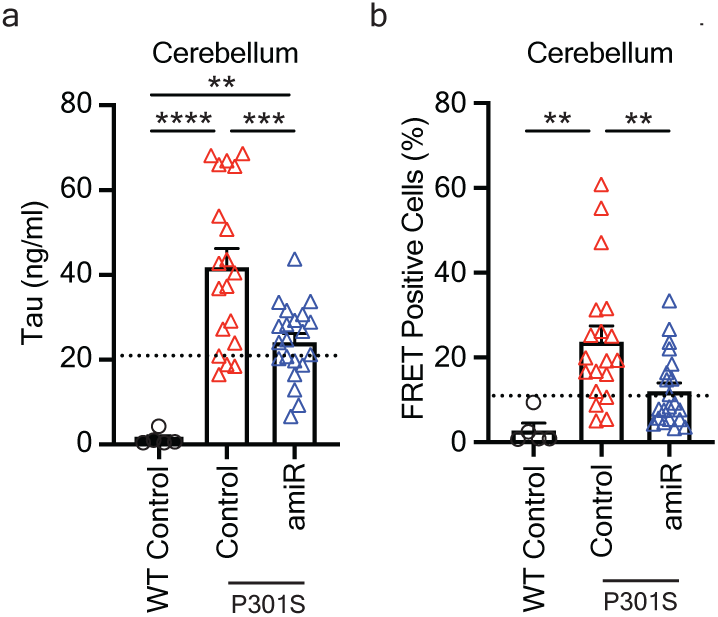
6-month efficacy data. **a** ELISA Quantification of 1N4R human P301S tau protein in the cerebellum of wildtype and P301S injected littermates; **b** Quantification of soluble tau seeding from the cerebellum. Mean±SD; One-way ANOVA with Tukey’s multiple comparisons; ** p< 0.01; ***p<0.001; ****p<0.0001. Dashed lines in histoplots represent the half maximal inhibitory value relative to the control group.

**Supplementary Figure 5.**
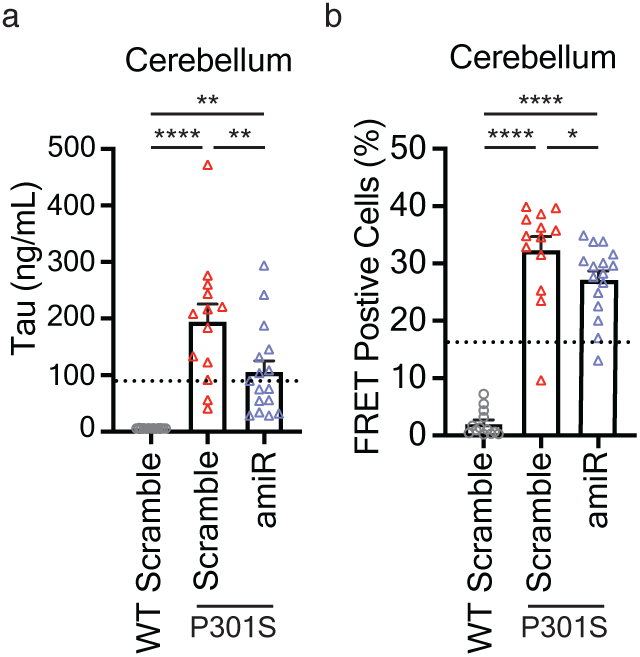
9-month efficacy data. **a,b** Histoplots illustrate tau protein **(a)** quantification by ELISA and tau seeding **(b)** from cerebellum of wildtype and P301S injected littermates. Mean±SEM; One-way ANOVA with Tukey’s multiple comparisons; * p< 0.05; ** p< 0.01; ****p<0.0001. Dashed lines in histoplots represent the half maximal inhibitory value relative to the control group.

**Supplementary Figure 6.**
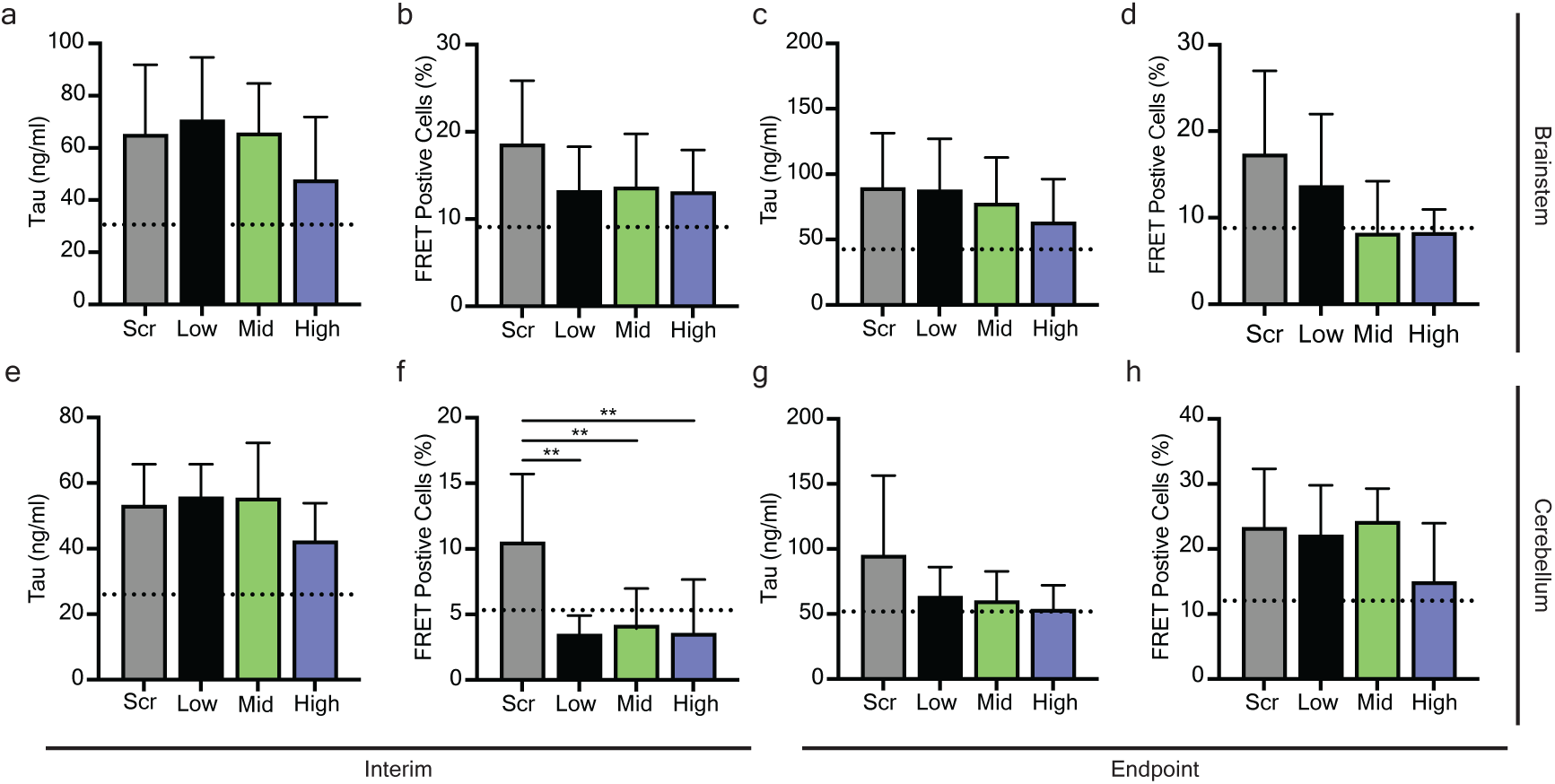
Dosing biochemical data. **a-h** Histoplots illustrate quantification of brainstem **(a-d)** and cerebellum **(e-h)** samples from interim **(a,b,e,f)** or endpoint **(c,d,g,h)** animal’s tau protein **(a,c,e,g)** and tau seeding **(b,d,f,h)**. Mean±SEM; One-way ANOVA with Tukey’s multiple comparisons; ** p< 0.01. Dashed lines in histoplots represent the half maximal inhibitory value relative to the control group.

