## Supplemental for "Preclinical Development of a Vectorized Artificial miRNA Gene Therapy for Tauopathies"

**Supplementary Figure 1. *In silico* prediction of artificial tau miRNAs. a,b** UNAFOLD hairpin 2 dimensional structures and sequences of lead human (**a**) and mouse (**b**) artificial miRNAs. Black and gray arrowheads indicate Drosha and Dicer cut sites respectively in the modified human miR-30a cassette and the underlined area indicates the guide miRNA strand sequence.

**Supplementary Figure 2. 3-month efficacy data.** **a** experimental paradigm illustrating three mo old WT or P301S mice injected intra-cisterna magna (ICM) with pre-clinical vector or vehicle control. **b** ELISA quantification of 1N4R human P301S tau cerebellum and cortex; **c** Soluble tau seeding of cerebellum and cortex lysates from wildtype and P301S littermates using HEK293T tau biosensors. Mean±SEM; One-way ANOVA with Tukey’s multiple comparisons; * p< 0.05; ** p< 0.01; ****p<0.0001. Dashed lines in histoplots represent the half maximal inhibitory value relative to the control group.

**Supplementary Figure 3. Preclinical and clinical vector designs display comparable total tau gene reduction.** **a** Illustration of experimental paradigm of the self-complimentary AAV constructs co-transfected into microwell plates with the MAPT dual luciferase screening constructs; **b,c** Histoplots illustrate the reduction of human 2N4R (**b**) and 2N3R (**c**) MAPT isoform knockdown using a dual luciferase reporter system. Mean±SD; One-way ANOVA with Tukey’s multiple comparisons; *** p< 0.001; ****p<0.0001. Dashed lines in histoplots represent the half maximal inhibitory value relative to the control group.

**Supplementary Figure 4. 6-month efficacy data.** **a** Experimental timeline following intra-cisterna magna (ICM) injection of 6 mo old WT and P301S mice. **b** ELISA Quantification of 1N4R human P301S tau protein in the cerebellum of wildtype and P301S injected littermates; **c** Quantification of soluble tau seeding from the cerebellum. Mean±SEM; One-way ANOVA with Tukey’s multiple comparisons; ** p< 0.01; ***p<0.001; ****p<0.0001. Dashed lines in histoplots represent the half maximal inhibitory value relative to the control group.

**Supplementary Figure 5. 9-month efficacy data.** **a** Experimental study design for testing treatment at late disease stage (9 mo of age). **b,c** Histoplots illustrate tau protein (**b**) quantification by ELISA and tau seeding (**c**) from cerebellum of wildtype and P301S injected littermates. Mean±SEM; One-way ANOVA with Tukey’s multiple comparisons; * p< 0.05; ** p< 0.01; ****p<0.0001. Dashed lines in histoplots represent the half maximal inhibitory value relative to the control group.

**Supplementary Figure 6. Dosing biochemical data.** **a** Experimental design for dosing 6 mo old P301S mice for interim (3 mo post-injection) and endpoint (6 mo post-injection) analysis groups. **b-i** Histoplots illustrate quantification of brainstem **(a-d)** and cerebellum **(e-h)** samples from interim **(b,c,d,e)** or endpoint **(f,g,h,i)** animal’s tau protein **(b,d,f,h)** and tau seeding **(c,e,g,i)**. Mean±SEM; One-way ANOVA with Tukey’s multiple comparisons; ** p< 0.01. Dashed lines in histoplots represent the half maximal inhibitory value relative to the control group.
